# Selection for or against Escape from Nonsense Mediated Decay is a Novel Signature for the Detection of Cancer Genes

**DOI:** 10.1101/2021.04.12.439236

**Authors:** Runjun D. Kumar, Briana A. Burns, Paul J. Vandeventer, Pamela N. Luna, Chad A. Shaw

## Abstract

Escape from nonsense mediated decay (NMD-) can produce activated or inactivated gene products, and bias in rates of escape can identify functionally important genes in germline disease. We hypothesized that the same would be true of cancer genes, and tested for NMD- bias within The Cancer Genome Atlas pan-cancer somatic mutation dataset. We identify 29 genes that show significantly elevated or suppressed rates of NMD-. This novel approach to cancer gene discovery reveals genes not previously cataloged as potentially tumorigenic, and identifies many potential driver mutations in known cancer genes for functional characterization.

---

Frameshift insertions, deletions and stop-gain point mutations usually generate new predicted termination codons (PTC+) and result in nonsense-mediated decay of the mRNA prior to translation (NMD+)^1^. However, termination codons within 55bp of the final exon-exon junction or further downstream can escape NMD (NMD-) and produce a truncated or extended peptide. Somatic NMD-variants are known to affect immunogenicity in some tumor types^2^, and the ratio of somatic PTC+ variants that are NMD+ or NMD- varies between tumor suppressors and oncogenes due to positive and negative selection on NMD+ variants^3^.

Studies in germline datasets suggest that nearly 10% of protein-coding genes may demonstrate selection on NMD- variants as well, particularly c-terminal events which produce truncated or extended peptides^4^. In some cases these events are thought to activate the peptide or produce a dominant-negative effect, and these mechanisms are important in known cancer genes^5, 6^. Identifying activating mutations in cancer is crucial as these events may be more easily targeted with small molecule inhibitors; however we have previously shown that existing methods struggle to identify these events^7^. We hypothesized that potentially activating c-terminal, PTC+ NMD- events would be enriched or depleted in some cancer genes in somatic cancer mutation data, and that new putative cancer genes could be identified with this novel mutation signature.

We first assessed whether PTC+ variants (stopgains and frameshift indels) have significantly higher or lower rates of NMD- versus PTC-variants (synonymous, missense and inframe indels) (Figure S1-2). NMD status is determined using the 55bp rule and the position of the predicted termination codon for PTC+ variants, and by the location of the variant itself for PTC- variants. Each gene was considered in the 0, +1 and −1 frames, with rates of NMD- compared between PTC- and PTC+ variants using Fisher exact tests in each frame. We then used Fisher’s method to combine p-values into a single result for each gene. After false-discovery-rate (FDR) correction, we identified 29 genes with abnormal rates of NMD-among PTC+ variants from 14,762 genes assessed (Table 1, Table S1). In 26 of these genes there was a higher rate of NMD- than expected; the remainder had relative depletions (Figure S3).

**Table 1.**
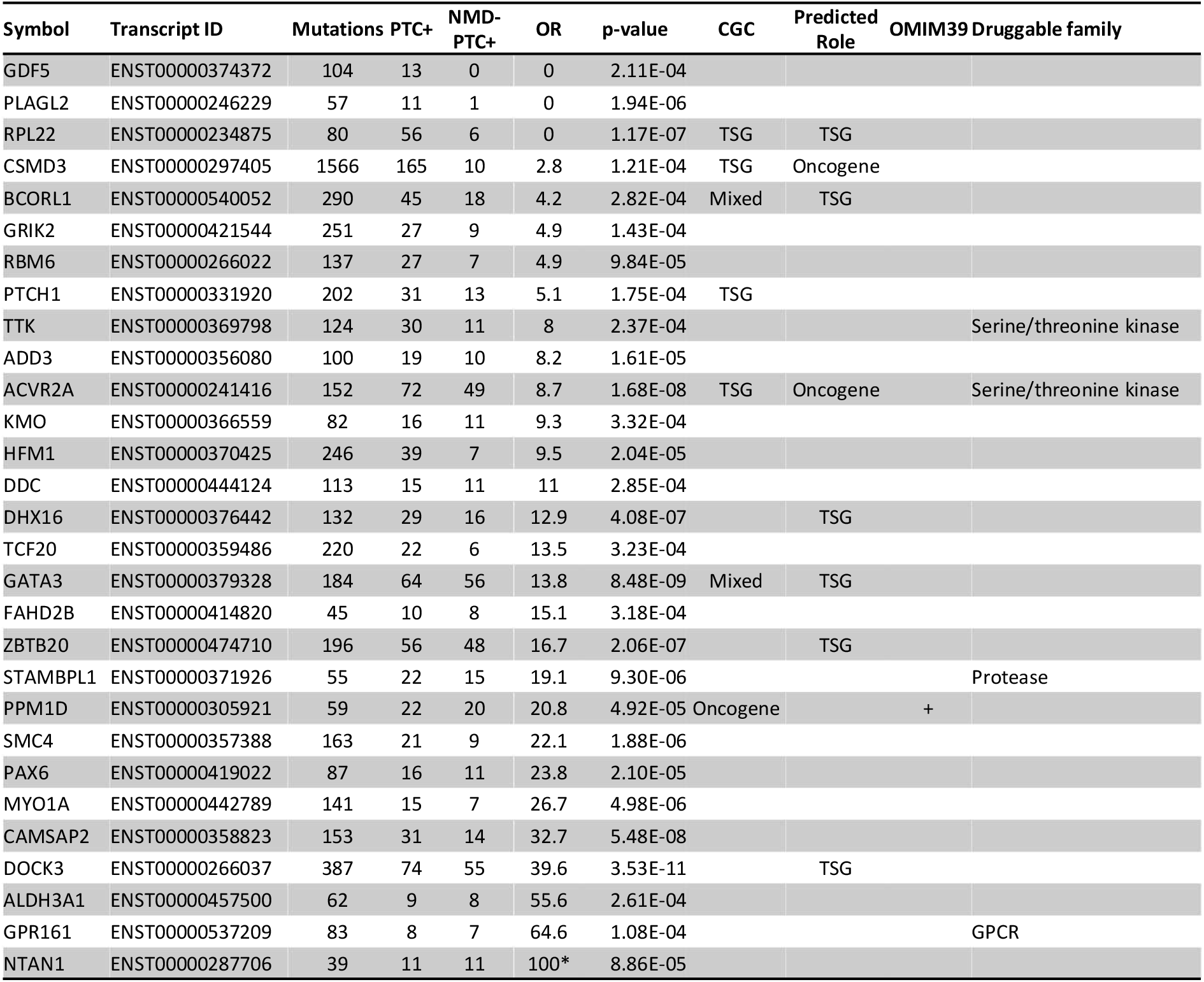
Genes with enrichment or depletion for NMD- among variants causing premature termination codons based on FDR < 0.2. Abbreviations: CGC, cancer gene census; GPCR, g-protein coupled receptor; NMD-, escape from nonsense mediated decay; OMIM39, 39 genes with dominant NMD- events per Online Mendelian Inheritance in Man; OR, odds-ratio; PTC+, variant causing premature termination codon; TSG, tumor suppressor gene. Odds ratios with a zero count in the denominator are marked (*).

Seven of 29 genes are also among 630 genes in the Cancer Gene Census (CGC)^8^ (p=0.0001): four are identified as exclusively TSGs (*ACVR2A, CSMD3, PTCH1, RPL22*), one exclusively as an oncogene (*PPM1D*), and two are annotated as having a mixed role (*BCORL1, GATA3*). We ran our previously reported ensemble method for cancer gene discovery as well (Figure S4), and it identified 3 additional genes as predicted cancer genes (*DHX16, DOCK3*, and *ZBTB20* as TSGs); the model also would classify *GATA3* and *BCORL1* as TSGs and *ACVR2A* and *CSMD3* as oncogenes^9^. Four genes were potentially druggable based on their gene family (*ACVR2A, GPR161, STAMBPL1* and *TTK*)^10^.

Interestingly, none of the 29 genes were identified in a prior analysis of NMD- depletion based on population germline datasets (Figure S5)^4^. However, *PPM1D* is one of 39 genes identified in the Online Mendelian Inheritance in Man (OMIM) as bearing activating c-terminal NMD- PTC+ variants, and these events are implicated in tumor development^6^. In general, we found evidence of NMD- bias in sets of cancer genes, but not in gene sets known to show NMD- bias in germline data sets (Figure S6).

Therefore, we assessed NMD- bias while limiting scope to sets of known cancer genes. We started with 87 high-confidence cancer genes identified prior to widespread tumor sequencing which we have previously used for benchmarking (“HiConf cancer genes”)^9^. Only 5/87 were significantly biased at a FDR of 0.2 (*GATA3, PTCH1, VHL, FBXW7, APC*, Figure 1A). There is a general skew among p-values for these genes compared with others, suggesting we may not be sufficiently powered to detect cancer genes with significant NMD- bias (Figure 1B, Figure S6). The identified genes had a variety of effect sizes (Figure 1C-G), and extending this analysis to 431 CGC oncogenes and TSGs additionally identified *ACVR2A, ASXL1, ASXL2, CASP8, CSMD3* and *PPM1D* with significant deviations at FDR<0.2 (Figure S7). We also analyzed nonsense variants in the COSMIC dataset^8^, and found that HiConf TSGs showed a relative depletion in NMD- compared with HiConf oncogenes (p=0.012, Figure S8), similar to the pattern in TCGA data, with the caveat that the COSMIC analysis contains only in-frame variants and combines different sequencing platforms.

**Figure 1.**
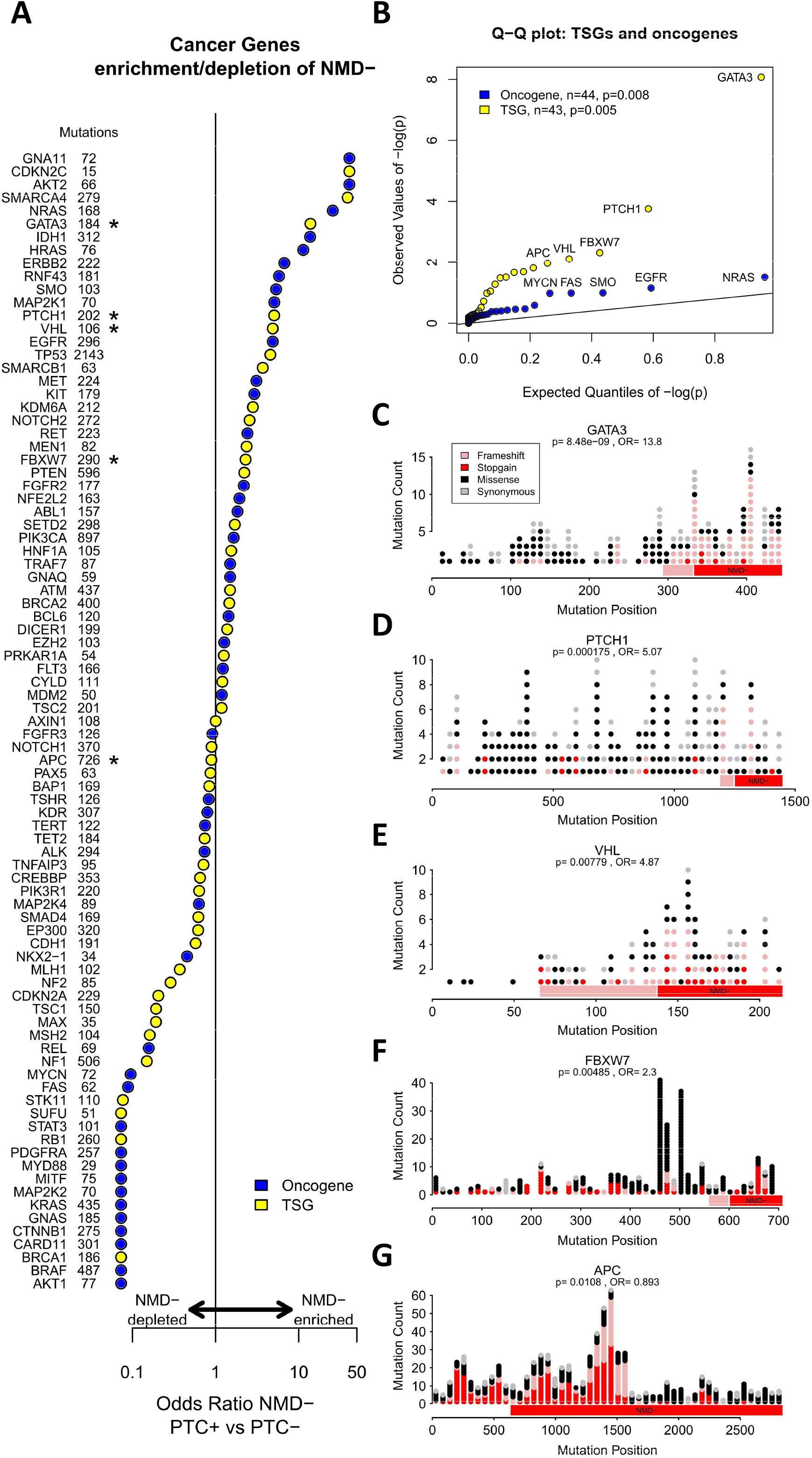
Enrichment or depletion of NMD- PTC+ events among oncogenes and tumor suppressors. A) Odds-ratios of NMD- for PTC+ compared with PTC-variants are shown for 87 high-confidence cancer genes. Mutation counts are displayed. Genes identified at FDR < 0.2 based on Fisher testing marked with (*). B) Q-Q plot of p-values for oncogenes and TSGs. C-G) Mutation diagrams for the 5 genes with FDR < 0.2. NMD- regions are labeled. The red area is NMD- in all frames. The pink area is NMD-for frameshift events. The boundary is drawn at the more extensive of −1 or +1 frames.

Analysis of NMD- bias offers insight into specific cancer genes. *PPM1D* represents one pattern we expect to find for an activating variant, with enrichment of c-terminal PTC+ NMD- events in a known oncogene^6^. *GATA3* is another example; it has a complex role in breast cancer, but there is evidence that some c-terminal frameshifts result in an extended transcript that has gain-of-function properties, while others shorten the transcript and have a dominant negative effect^11^. The results of this analysis may have similar implications for the other known cancer genes identified in Table 1, several of which are usually categorized as tumor suppressors. PTC+ NMD- variants in these genes may cause loss-of-function despite expression of the peptide, which could potentially be rescued pharmacologically, or they may alter the function of the peptide in some other way, expanding the role of the gene in tumorigenesis.

Our results also implicate several genes in cancer development for the first time. We identify that *growth/differentiation factor 5* (*GDF5*) is deficient in NMD- PTC+ variants, as would be expected of a tumor suppressor for the gain-of-function mechanism of NMD- PTC+ variants; however in this case the absolute number of PTC+ variants is small, and the role of *GDF5* in tumorigenesis is not well established^12-14^. Many of the remaining genes in Table 1 have recurrent NMD- PTC+ variants that could produce activated and truncated peptides. For example, there is a recurrent event K405Rfs*21 event in *STAM-binding protein-like 1* (*STAMBPL1*). This gene is not well understood, though it appears to be positively associated with prostate and gastric cancers^15, 16^, and mutations in this region have been implicated in canine polyp development^17^.

The use of NMD- bias among PTC+ variants is a promising new strategy for identifying cancer genes which is substantially orthogonal to prior methods. While the effect on protein function may vary across NMD- variants, the inference that PTC+ variants cause loss-of-function does not hold in the NMD- case, and these variants present a novel class for oncological analysis. We developed a framework for identifying NMD- patterns of association within cancer somatic mutation datasets. We identified 29 genes that demonstrated significant bias for PTC+ in the NMD- region. The majority were enriched in NMD- events which could represent activating c-terminal truncations. Many of these genes are potential cancer genes. These variants also represent novel targets in several established cancer genes, many of which are currently classified as TSGs. We anticipate that this strategy could be even more fruitful in larger datasets, which would also allow further analysis within specific cancer sub-types.

## Acknowledgements

We thank Dr. Zeynep Coban Akdemir for critically reading the manuscript and providing helpful feedback. We thank Dr. William Decker and his laboratory for wet lab mentorship of B.A. B., as well as funding her work through Alex’s Lemonade Stand.

## Methods

### Datasets

Pan-cancer TCGA annotated mutations were downloaded from the public repository (https://gdc.cancer.gov/about-data/publications/mc3-2017, filename mc3.v0.2.8.PUBLIC.maf.gz, accessed 12/20/2020). Cancer Gene Census tier 1 and tier 2 genes were downloaded from COSMIC (https://cancer.sanger.ac.uk/census, accessed 12/20/2020). Other gene sets or annotations are drawn directly from the cited works unless specifically noted. We used the GenomOncology Precision Oncology API Suite (https://www.genomoncology.com/api-suite) to identify the distance of each variant from the exonic boundaries and determine NMD status.

### Annotations and Filtering

Annotations are directly from the TCGA MAF file whenever possible. Only coding mutations (stopgains, frameshift indels, synonymous variants, missense variants, and inframe indels) were considered in the analysis. Variants were annotated with the frame they would produce. By definition, stopgains, synonymous, missense and inframe indels do not change the frame and are assigned a frame of 0. Frameshift indels that result in the frame moving forward by one position (e.g. deletion of one, four, seven base pairs, or the insertion of two, five, eight base pairs, *etc*) are assigned frame +1. Frameshift indels that result in the frame moving backward by one position are assigned frame −1.

We annotated mutations as being PTC+ (frameshift indel, stop-gains) or PTC- (synonymous, missense, inframe indel) and then annotated the PTC+ variants as being NMD+ or NMD- based the position of the predicted termination codon and the 55bp rule as previously described^4^. Each transcript was divided into NMD- and NMD+ regions for each potential reading frame. The boundary is defined as the most upstream position where a variant of that frame would produce a PTC within 55bp of the final exon-exon junction or further downstream. The transcript used was the canonical transcript used in the original TCGA dataset. There were 1,306,510 mutations affecting 16,037 genes in 9,859 individuals with 34 cancer types at this point.

### Analysis

We next confirmed the most appropriate control to assess rates of NMD- among PTC+ variants. We showed that synonymous mutations do not occur randomly between NMD+ and NMD- regions of the gene in at least some genes, and therefore chose observed mutation rates as a control (Figure S1). Synonymous mutations make up roughly one third of point mutations in the dataset; to maximize data usage, we assessed whether missense mutations were comparable to synonymous events with regards to occurrence within the NMD- region (Figure S2). We could find no genes where missense and synonymous variants significantly differed with regards to occurrence within the NMD- region. Therefore pooled synonymous and missense variants (PTC- variants) were compared to PTC+ variants to assess rates of NMD- (Figure S3). These control experiments did not test the +1 or −1 frames since missense and synonymous mutations occur exclusively in-frame.

We tested for NMD- bias by calculating a Fisher exact test for 2 by 2 tables for each frame in each gene (Figure S3). The p-values from these tests were combined using the Fisher Method, resulting in a single p-value for each gene. A pooled odds-ratio was also calculated by combining the counts across the three frames for each gene. These pooled odds-ratios and combined p-values are used throughout the manuscript unless otherwise noted.

Fisher Exact Tests were also used to assess the overlap between different gene sets throughout the analysis. False Discover Rates were controlled using Benjamini-Hochberg correction. To compare sets of genes, chi-square statistics were calculated for each frame and gene in the set and summed; this value was then compared to random draws of genes to create p-values. Methods and techniques for comparison analyses (e.g. druggability, cancer gene prediction) are found in the pertinent references.

---

**Figure S1.**
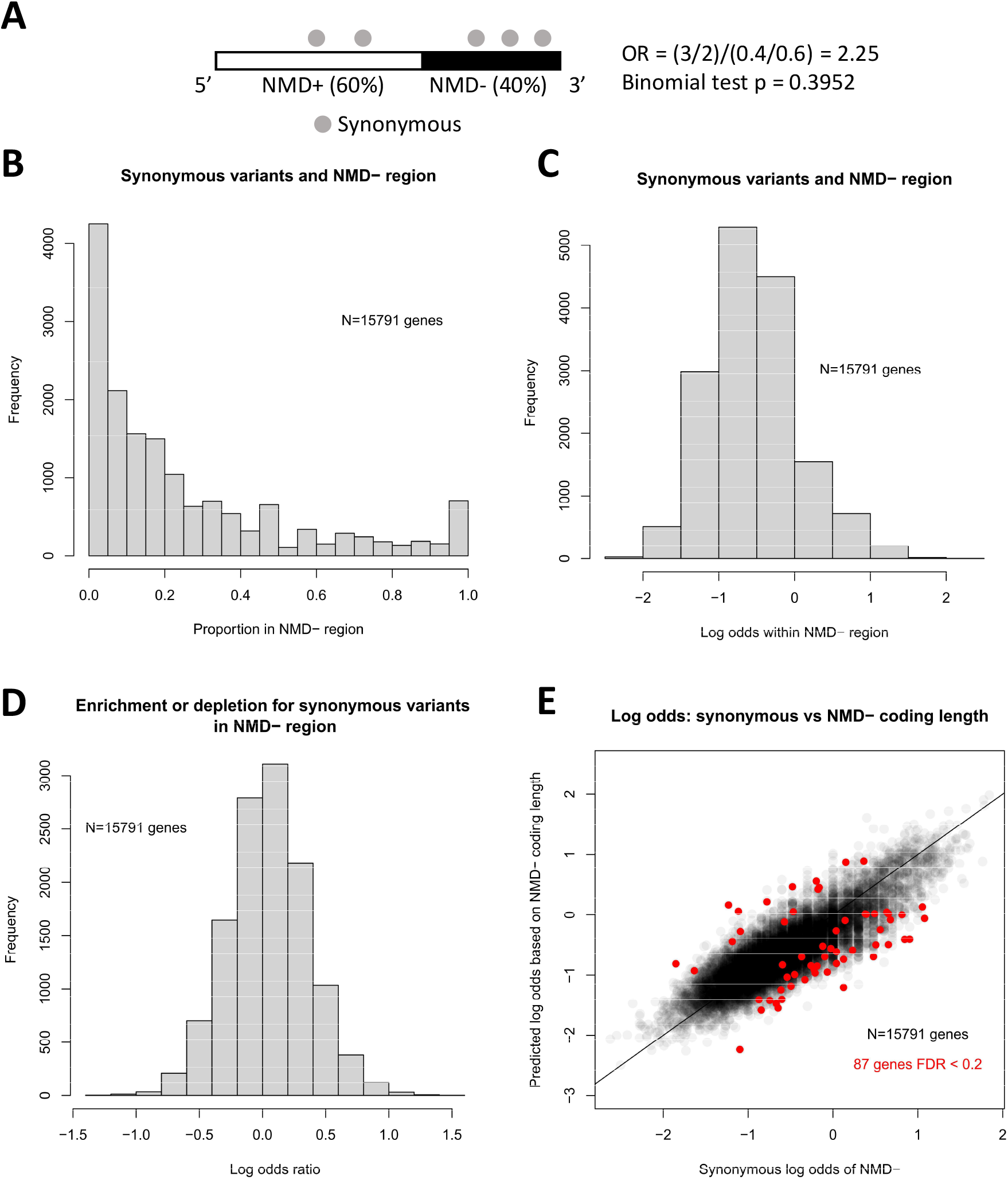
Synonymous variants do not occur randomly between NMD+ and NMD- regions. A) Schema for the analysis. A mature transcript is depicted with an NMD- region that is 40% of the coding length. B) Proportion of synonymous variants in the NMD- region for each gene. −1 and +1 frames are excluded. C) Log odds of synonymous variants occurring in the NMD-region. D) Log odds-ratios for synonymous variants occurring in the NMD- region, calculated as outlined in panel (A). E) Genes plotted by observed and expected odds of NMD- among synonymous variants, assuming variants occur randomly throughout the coding region. 87 genes with significant differences are highlighted.

**Figure S2.**
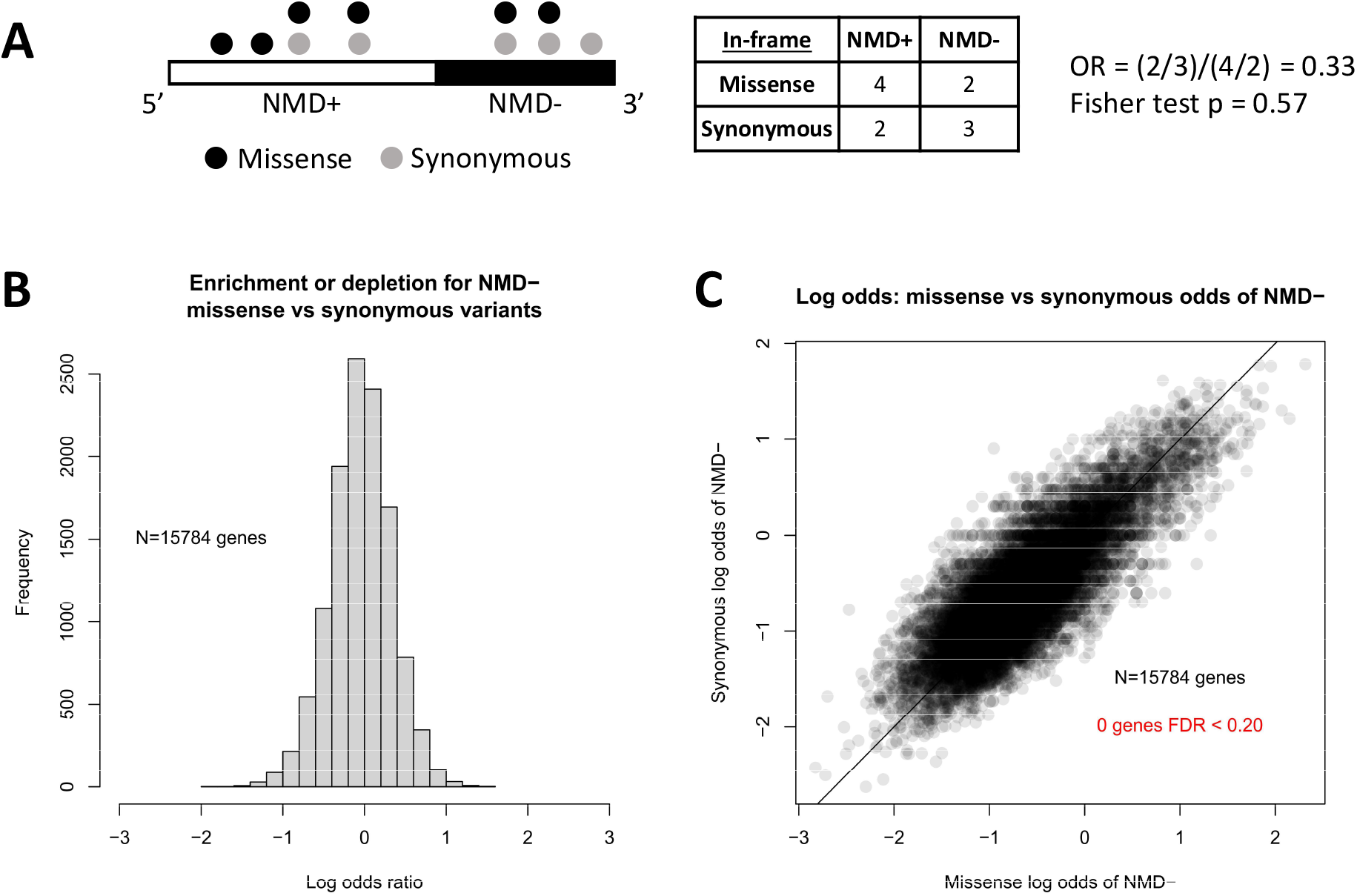
Missense and synonymous variants do not significantly differ with regards to presence in NMD- regions. A) Schema of analysis. A mature transcript is depicted. B) Log odds-ratios for rate of missense variants in NMD- region versus synonymous variants, as outlined in panel (A). −1 and +1 frames are excluded. C) Genes plotted by observed log odds of NMD- for missense and synonymous variants. The rates are compared with Fisher exact testing with no genes showing significant differences.

**Figure S3.**
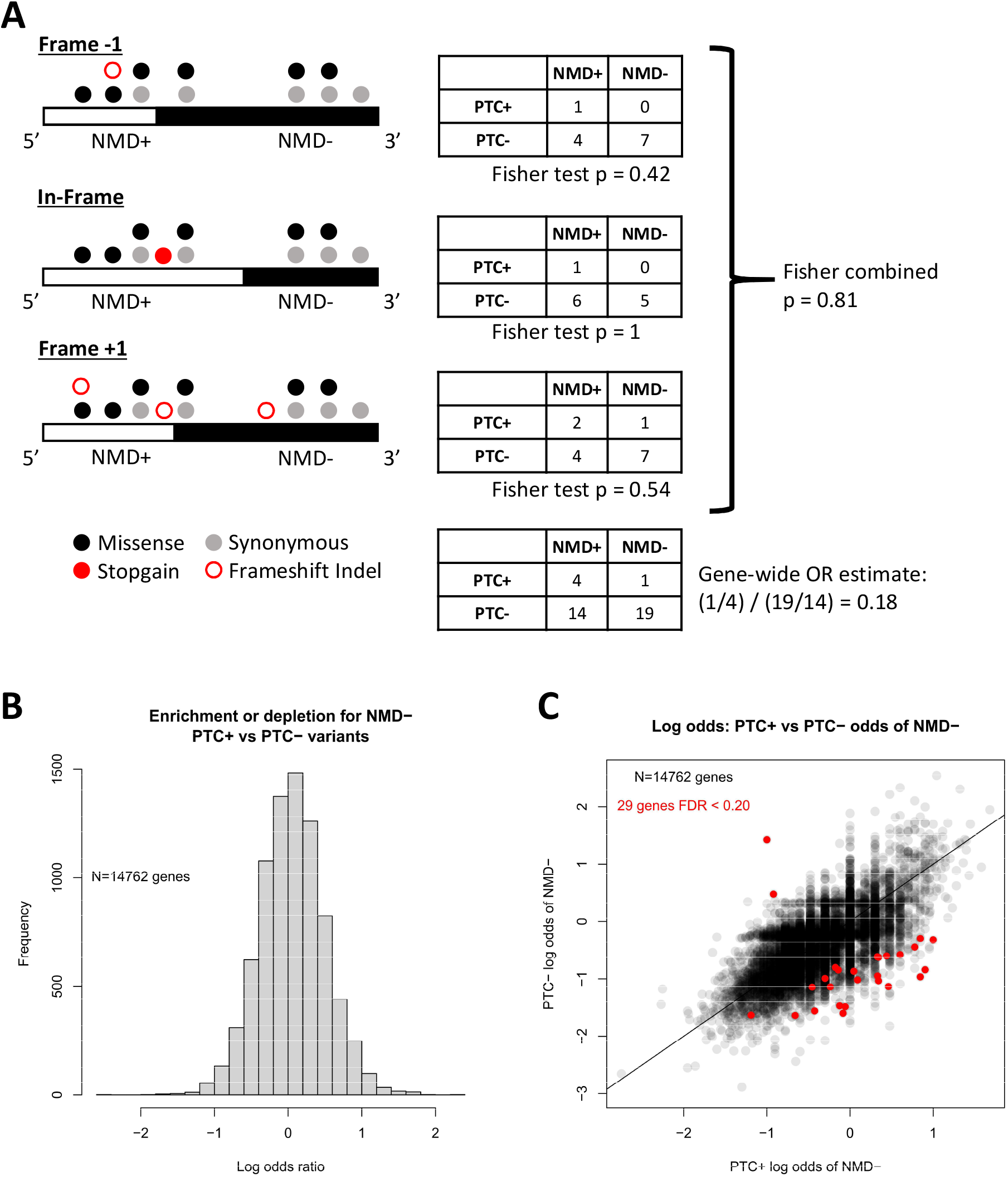
Analysis of PTC+ variants versus PTC-variants with regards to presence in NMD-region. A) Schema of analysis. A mature transcript is depicted, with NMD- regions defined for each reading frame using the 55bp rule. B) Log odds-ratios for rate of PTC+ variants in NMD- region versus synonymous and missense (PTC-) variants, as outlined in panel (A). C) Genes plotted by observed log odds of NMD- for PTC+ and PTC-variants. The 29 genes with significant NMD- bias are highlighted.

**Figure S4.**
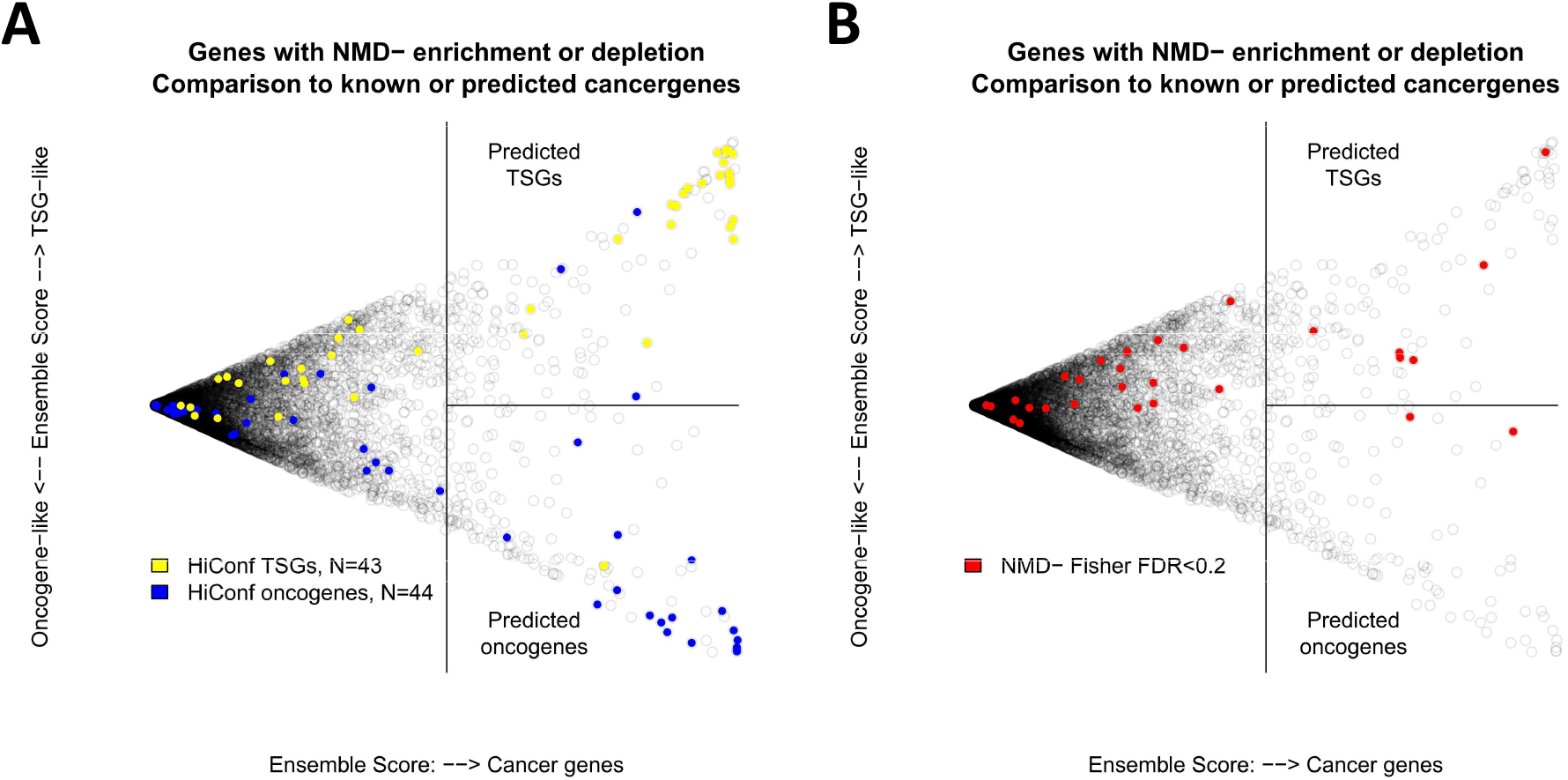
Prediction of tumor suppressors and oncogenes and comparison to genes enriched or depleted in NMD- PTC+ variants. A) An ensemble method identifies predicted oncogenes (lower right segment) and predicted TSGs (upper right segment). High confidence cancer genes identified prior to widespread tumor sequencing are highlighted (“HiConf cancer genes”). B) The same plot with the 29 genes showing NMD- bias highlighted.

**Figure S5.**
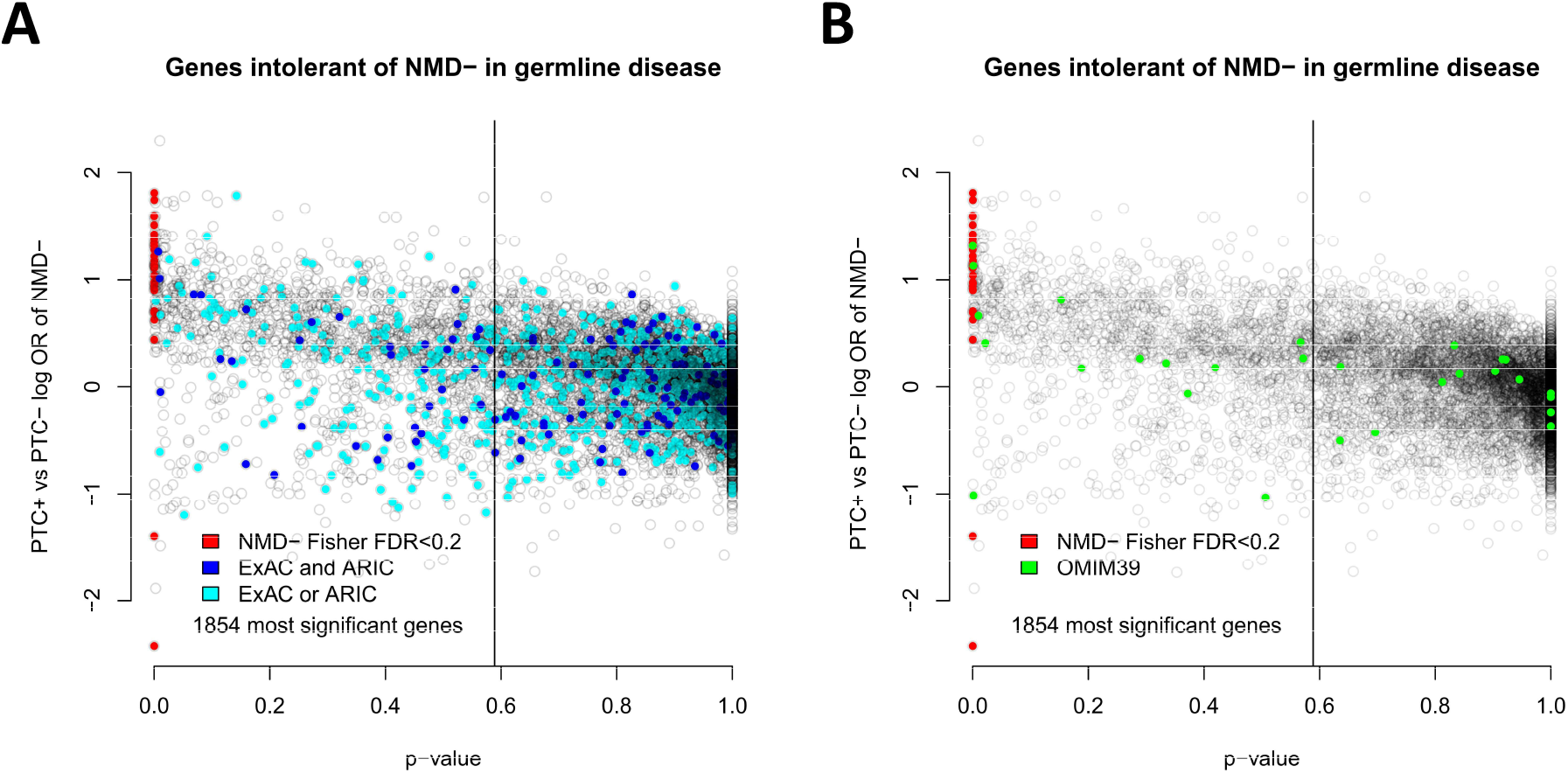
Comparison of current analysis in somatic cancer datasets to genes identified previously in germline datasets. “ExAC and ARIC” genes were identified in both of those two datasets as being depleted in NMD- PTC+ variants. “ ExAC or ARIC” genes were identified in one dataset or the other. OMIM39 genes are annotated in OMIM as having NMD- PTC+ variants that cause dominant disease. To the left of the vertical line are as many genes as identified in the ExAC or ARIC set (n=1854).

**Figure S6.**
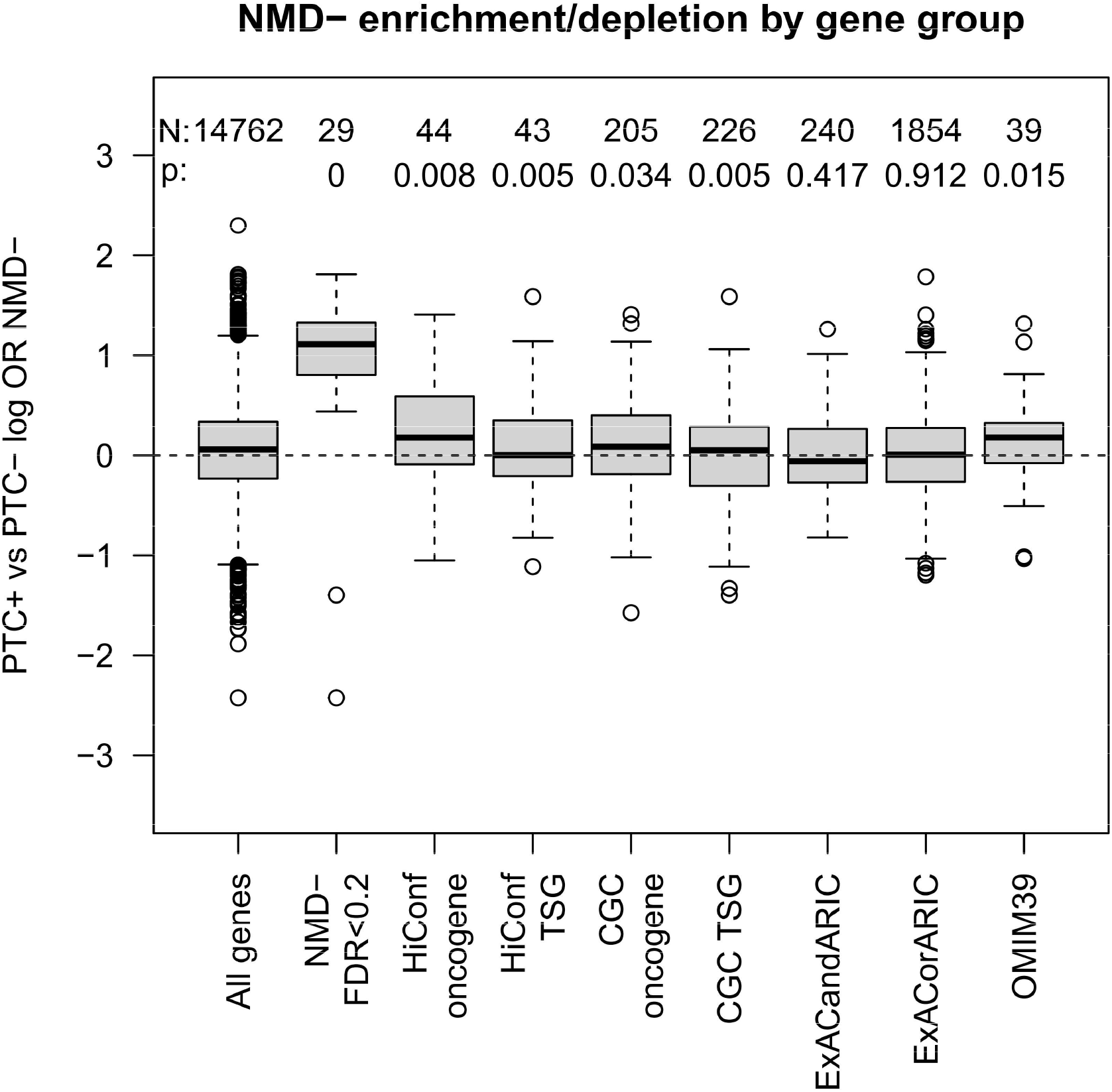
Log odds-ratios for NMD- enrichment or depletion among PTC+ variants by gene set. p-values indicate chi-square testing for the class as a whole, and test for abnormal enrichment or depletion among members of the class versus a randomly drawn set of equal size.

**Figure S7.**
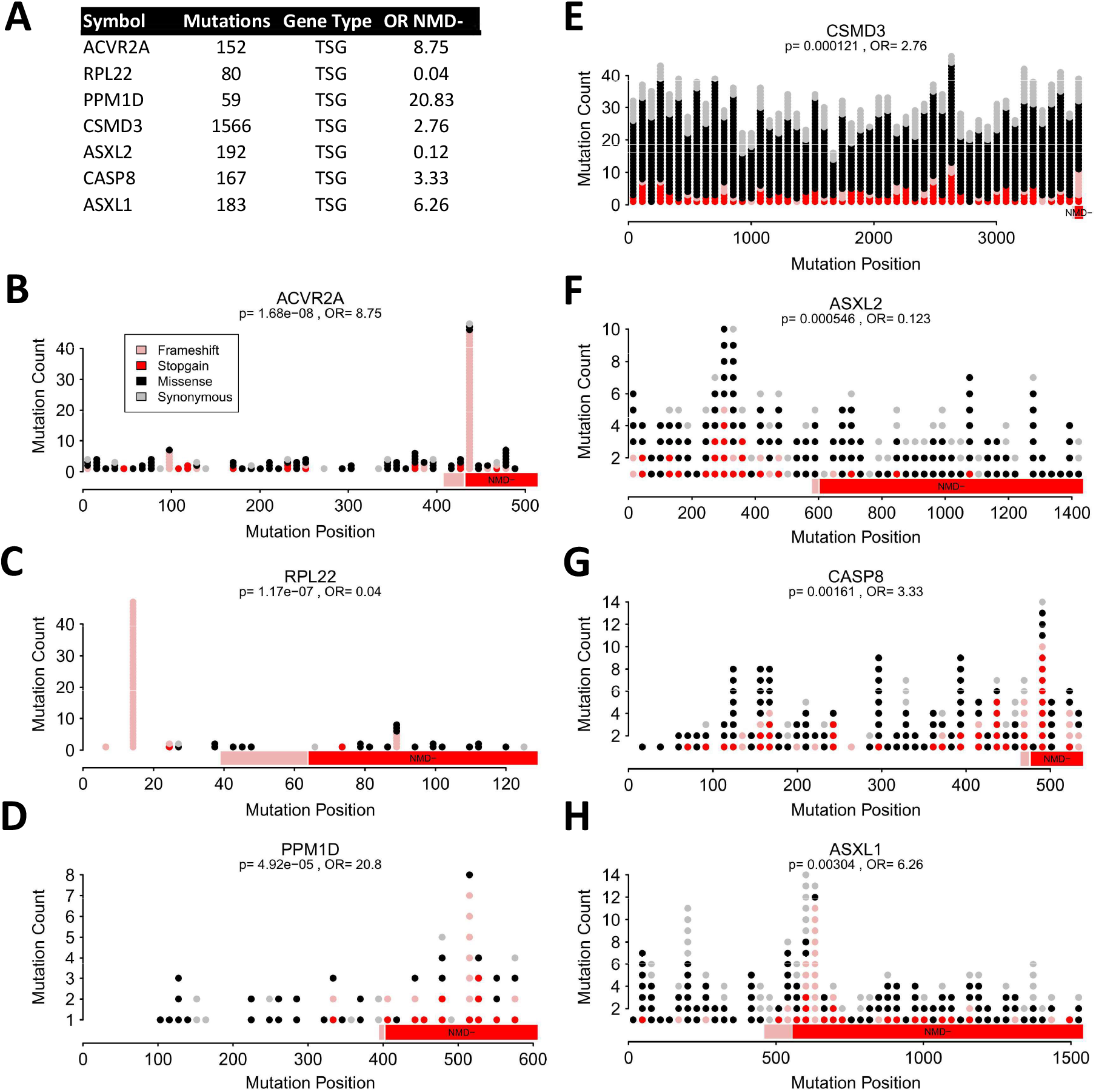
Additional oncogenes and tumor suppressor genes from the Cancer Gene Census with significant NMD- bias.

**Figure S8.**
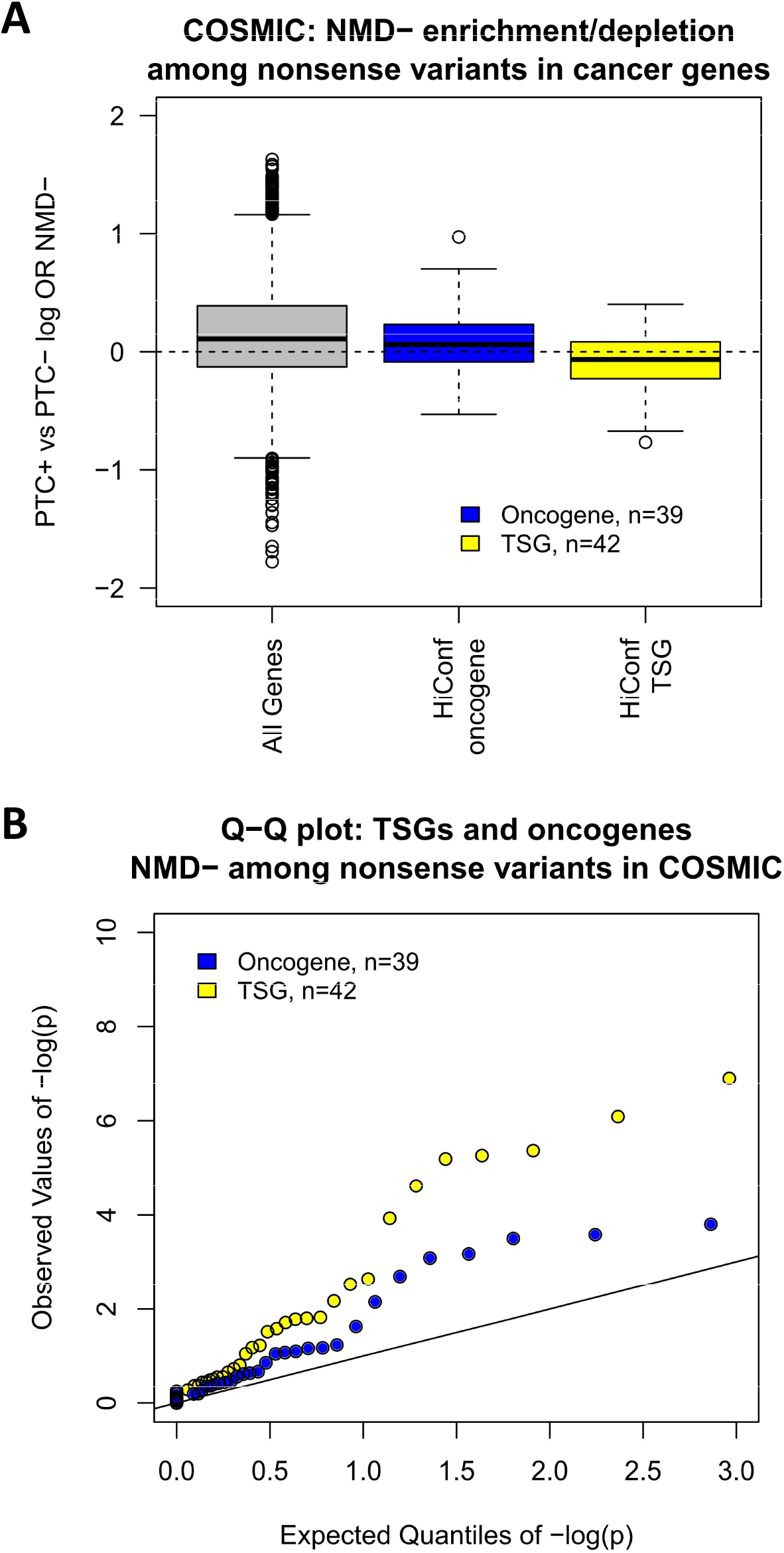
High-confidence cancer genes show similar patterns of NMD- among nonsense variants in the COSMIC dataset. A) Boxplot of log odds-ratios for all genes, HiConf oncogenes, and HiConf tumor suppressors based on nonsense variants only (frame=0) in the COSMIC dataset. Tumor suppressors are relatively NMD- depleted relative to the oncogenes (Student’s t-test: p=0.012). B) QQ plot of p-values for the high confidence oncogenes and tumor suppressors. Two outliers with observed -log(p) > 15 are omitted.

